# Pseudoneglect in visual search: Behavioral evidence and connectional constraints in simulated neural circuitry

**DOI:** 10.1101/129171

**Authors:** Onofrio Gigliotta, Tal Seidel Malkinson, Orazio Miglino, Paolo Bartolomeo

## Abstract

Most people tend to bisect horizontal lines slightly to the left of their true center (pseudoneglect), and start visual search from left-sided items. This physiological leftward spatial bias may depend on hemispheric asymmetries in the organization of attentional networks, but the precise mechanisms are unknown. Here we modeled relevant aspects of the ventral and dorsal attentional networks (VAN and DAN) of the human brain. First, we demonstrated pseudoneglect in visual search in 101 right-handed psychology students. Participants consistently tended to start the task from a left-sided item, thus showing pseudoneglect. Second, we trained populations of simulated neurorobots to perform a similar task, by using a genetic algorithm. The neurorobots’ behavior was controlled by artificial neural networks, which simulated the human VAN and DAN in the two brain hemispheres. Neurorobots differed in the connectional constraints that were applied to the anatomy and function of the attention networks. Results indicated that (1) neurorobots provided with a biologically plausible hemispheric asymmetry of the VAN-DAN connections, as well as with inter-hemispheric inhibition, displayed the best match with human data; however, (2) anatomical asymmetry *per se* was not sufficient to generate pseudoneglect; in addition, the VAN must have an excitatory influence on the ipsilateral DAN; (3) neurorobots provided with bilateral competence in the VAN but without inter-hemispheric inhibition failed to display pseudoneglect. These findings provide a proof of concept of the causal link between connectional asymmetries and pseudoneglect, and specify important biological constraints that result in physiological asymmetries of human behavior.

**Significance statement:** Most of us start our exploration of the environment from the left side. Here we demonstrated this tendency in undergraduate students, and trained artificial agents (neurorobots) to perform a similar visual search task. The neurorobots’ behavior was controlled by artificial neural networks, inspired by the human fronto-parietal attentional system. In seven distinct populations of neurorobots, different constraints were applied on the network connections within and between the brain hemispheres. Only one of the artificial populations behaved in a similar way to the human participants. The connectional constraints applied to this population included known characteristics of the human fronto-parietal networks, but had also additional properties not previously described. Thus, our findings specify biological constraints that induce physiological asymmetries of human behavior.

## 1. Introduction

A thorough exploration of the space around us is essential to everyday life. However, spatial exploration is not perfectly symmetrical in humans. For example, when we explore a scene in order to cancel out visual targets, we tend to start the search from the left part of the sheet (Azouvi et al., 2006; Bartolomeo, D’Erme, & Gainotti, 1994). This physiological leftward spatial bias is analogous to the slight physiological leftward shift typically observed in horizontal line bisection, termed pseudoneglect (Bowers & Heilman, 1980) because it goes in the opposite direction to the typical rightward bias showed by patients with left visual neglect after right hemisphere damage (Schenkenberg, Bradford, & Ajax, 1980; Urbanski & Bartolomeo, 2008).

Evidence shows that visuospatial attention is a major determinant of pseudoneglect (McCourt, Garlinghouse, & Reuter-Lorenz, 2005; Toba, Cavanagh, & Bartolomeo, 2011), which might thus result from asymmetries in the hemispheric control of attention (McCourt & Jewell, 1999; Ossandón, Onat, & König, 2014). However, the specific neural structures and the mechanisms at the basis of pseudoneglect remain unknown.

In the human brain, visuospatial attention is controlled by fronto-parietal networks, which demonstrate substantial asymmetries favoring the right hemisphere (Corbetta & Shulman, 2002; Heilman & Van Den Abell, 1980; Mesulam, 1999). Dysfunction of these networks after right hemisphere damage can induce signs of neglect for left-sided events (Bartolomeo, Thiebaut de Schotten, & Chica, 2012; Corbetta & Shulman, 2011). A bilateral dorsal attentional network (DAN), composed by the intraparietal sulcus / superior parietal lobule and the frontal eye field / dorsolateral prefrontal cortex, shows increased BOLD responses during the orienting period (Corbetta & Shulman, 2002). A right-lateralized ventral attentional network (VAN) includes the temporoparietal junction and the ventrolateral prefrontal cortex. The VAN is important for detecting unexpected but behaviorally relevant events, and induces the DANs to reorient attention towards these events. Anatomically, three branches of a long-range white matter pathway, the Superior Longitudinal Fasciculus (SLF), connect these networks. The SLF has a ventro-dorsal gradient of hemispheric asymmetry (Thiebaut de Schotten et al., 2011). The ventral branch (SLF III) connects the VAN and is anatomically larger in the right hemisphere than in the left hemisphere, whereas the dorsal branch (SLF I, connecting the DAN) is more symmetrical. The lateralization of the intermediate branch (SLF II) displays interindividual differences, and is strongly correlated to the individual amount of pseudoneglect in line bisection and to differences in the speed of detection between left-sided and right-sided targets. Specifically, larger SLF volumes in the right hemisphere correlate with larger leftward bias (Thiebaut de Schotten et al., 2011).

A further potential source of performance asymmetry resides in the pattern of inter-hemispheric connections. Behavioral and electrophysiological evidence suggests that inter-hemispheric communication is not strictly symmetrical in humans, but it is faster from the right to the left hemisphere (Marzi, 2010). Also, the posterior callosal connections from the right parietal node of the DAN to its left hemisphere homologue seem to be predominantly inhibitory (Koch et al., 2011). Concerning the VAN, its right and left temporo-parietal caudal nodes are not strongly connected by callosal fibers (Catani & Thiebaut de Schotten, 2012), and thus work in relative isolation from one another.

It is tempting to relate these biological constraints to the widespread leftward bias that occurs in human exploratory behavior. However, little is known about the specific dynamic interplay between the attentional networks resulting in pseudoneglect. On the one hand, methods used in humans have substantial limitations of spatiotemporal resolution and of inferential power, which severely limit their scope. On the other hand, it is difficult to draw firm conclusions from monkey neurophysiology, because of important differences between humans and primates in the organization of attention networks (Patel et al., 2015). In the present study, we took a different approach to unravel these issues. First, we tested a group of human participants to establish the presence and characteristics of pseudoneglect in a visual search task (Experiment 1). In Experiment 2, we trained neurally controlled robots (neurorobots) to perform a task as similar as possible to the human one. We then articulated detailed implementations of several instances of attention network architecture, which directed the neurorobots’ performance, in order to identify the structural and functional network constraints crucial for simulating human performance.

## 2. Experiment 1: Pseudoneglect in human visual search

### 2.1 Introduction

Pseudoneglect has been mainly measured using tasks of perceptual estimation of the length of horizontal lines (Bowers & Heilman, 1980; Jewell & McCourt, 2000; Toba et al., 2011). Analogous leftward biases seem also to occur in visual search tasks, as a tendency to find first targets on the left side of the display (Azouvi et al., 2006; Bartolomeo et al., 1994), but evidence in this domain is much less systematic. Thus, in the present context it was important to test our specific task in order to ensure the validity of the human-robotic comparison of performance.

### 2.2. Methods

#### 2.2.1. Ethics Statement

The procedure was approved by the local ethics committee.

#### 2.2.2 Participants

A total of 101 right-handed psychology students (76 females; mean age ± SD, 22.24 ± 4.40) gave their informed consent to perform a visual search experiment for course credit.

#### 2.2.3. Procedure

The task was designed to be as close as possible to that performed by neurorobots (see section 3 below). Participants were instructed to cancel as fast as possible targets displayed on a touch-sensitive tablet (Mediacom Winpad 801 8-inches, 120 dpi, 1280x800 pixels, refresh frequency 60 Hz), by using a stylus pen. Participants were comfortably seated with a viewing distance of ~40 cm. Each session consisted of 30 trials. Each trial was initiated by the participant touching a green round button placed at the center of the screen. Subsequently, a set of 5 dark-red (HEX #800000) filled round targets, with a 40-pixel radius (0.76° visual angle), was presented. Targets were randomly scattered on a display area of 512x512 pixels (9.7° × 9.7°), placed at the center of the screen. Upon participant’s touch, cancelled targets became bright red (HEX #FF0000). To assess lateral bias, we first defined the center of the display as 0, so that the values of the X coordinate went from -256 pixels (-4.85°) on the extreme left to +256 pixels (+4.85°) on the extreme right. Second, we measured the average position on the X axis of the first cancelled stimulus for each trial.

### 2.3. Results

As expected with this easy task, accuracy was at ceiling, with all participants correctly cancelling all the targets. Results showed a left-biased distribution of the first found target (see Fig. 9A below). The average X value was -80.23 pixels (1.52°), which significantly differs from the central position at X = 0 (Wilcoxon-Mann-Whitney two-tailed test, Z=-6.37, *p*<0.001).

### 2.4. Discussion

During a visual search task similar to that used for our simulations, normal participants exhibited a leftward bias (pseudoneglect), consisting of a tendency to start the visual search from a left-sided target. This result was observed in an experimental setting as close as possible to that used for neurorobots, and replicates and extends previous results obtained with different types of visual search tasks, such as the line cancellation test (Bartolomeo et al., 1994) and the bells test (Rousseaux et al., 2001).

## 3. Experiment 2: Visual Search in Neurorobots

### 3.1. Introduction

A neurorobot is a real or simulated robot whose behavior is controlled by an artificial neural network. For the present experiment, we developed distinct populations of simulated neurorobots controlled by artificial neural networks with different connectivity constraints. The neurorobots’ task was designed to be as close as possible to that performed by human participants in Experiment 1.

### 3.2. Models

The simulated robot (Fig. 1) has a single artificial eye and an actuator (simulated hand) able to perform the cancellation task. The robot’s eye can move and zoom, and can thus be described as a pan/tilt/zoom camera, because it can move along the horizontal and vertical axes and can zoom in a range between 1x to 12x. The use of a zoom was inspired by models of attention, which stipulate that attention can either be distributed over the whole field, but with low resolving power, or be continuously constricted to small portions of the visual field with a concomitant increase in processing power (Eriksen & Yeh, 1985).

**Figure 1.**
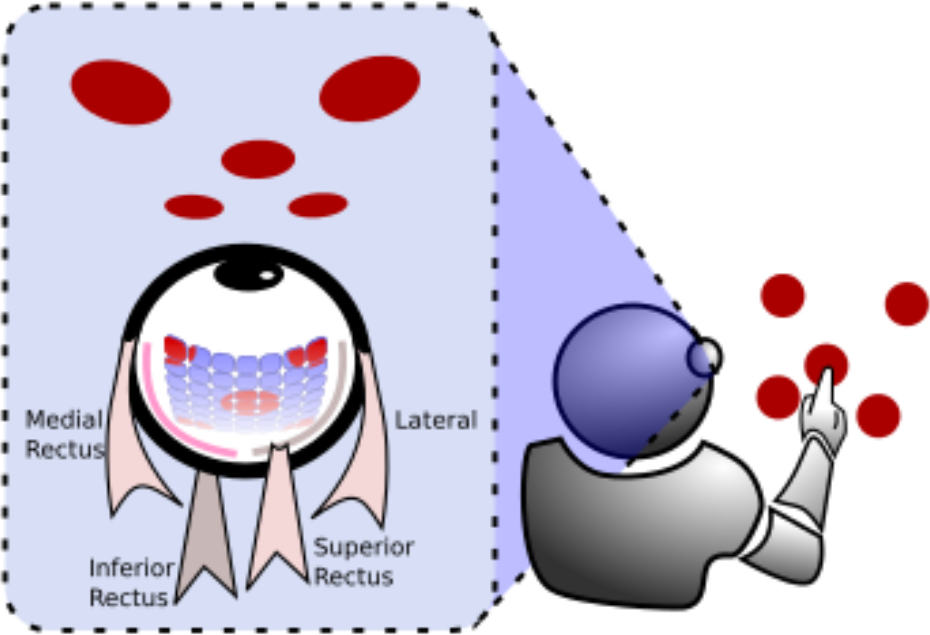
Schema of the neurorobot equipped with an artificial eye, provided with a 7×7 light receptor retina, and controlled by two pairs of simulated extraocular muscles.

The artificial eye is equipped with a retina made up of a 7×7 grid of light receptors (see Fig. 1). Each receptor outputs an activation value computed by averaging the luminance of the perceived stimuli across the receptive field, with radius set to 80 pixels. Receptors are evenly distributed within the artificial retina, which has a square form with a side varying from 1120 pixels (no zoom) to 96 pixels (maximum zoom). Thus, each stimulus can occupy a retinal surface ranging from 0.8% to 100% of the artificial retina. Horizontal and vertical movements of the eye are controlled by four simulated muscles (Massera, Ferrauto, Gigliotta, & Nolfi, 2014) (see Fig. 1), in analogy to the medial, lateral, inferior and superior recti of the human eye.

#### 3.2.1. Neural network

We used a standard neural network model in which each node of the network has a sigmoid activation function φ(x)=1/(1+e^-x^) and an adjustable threshold ϑ. The output, *O*, is computed for each node *i* by using the following equation:

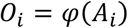

Where:

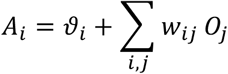

w_*ij*_ is the synaptic weight connecting unit *j* with unit *i*. The pattern of connections between nodes has been chosen according to biological evidence on dorsal and ventral attentional networks in human brains (see below, section 3.5).

Fig. 2A depicts the general template network. The 7×7 retina, consisting of 49 artificial neurons, constituted the input layer. The output layer controlled the zoom with two artificial neurons, the extraocular muscles with four neurons, and a decision unit for target detection, which triggered the touch response when exceeding a criterion threshold of 0.7. The hidden layer contained the attention networks and a hidden network devoted to control vertical eye movements (4 neurons, not depicted in Fig. 1). We modeled the DAN and the VAN by building a neural model organized across two hemispheres, with visual information from each visual field projecting to the contralateral hemisphere. Each DAN had 5 artificial neurons; each VAN had 4 artificial neurons. These parameters were based on pilot work, and reflect a tradeoff between network complexity and the time needed to run simulations. With these parameters, each simulation required about a week to be completed on our hardware. The VAN-DAN connections in the right hemisphere outnumbered those in the left hemisphere, in order to simulate analogous results for the human SLF II (Thiebaut de Schotten et al., 2011).

**Figure 2.**
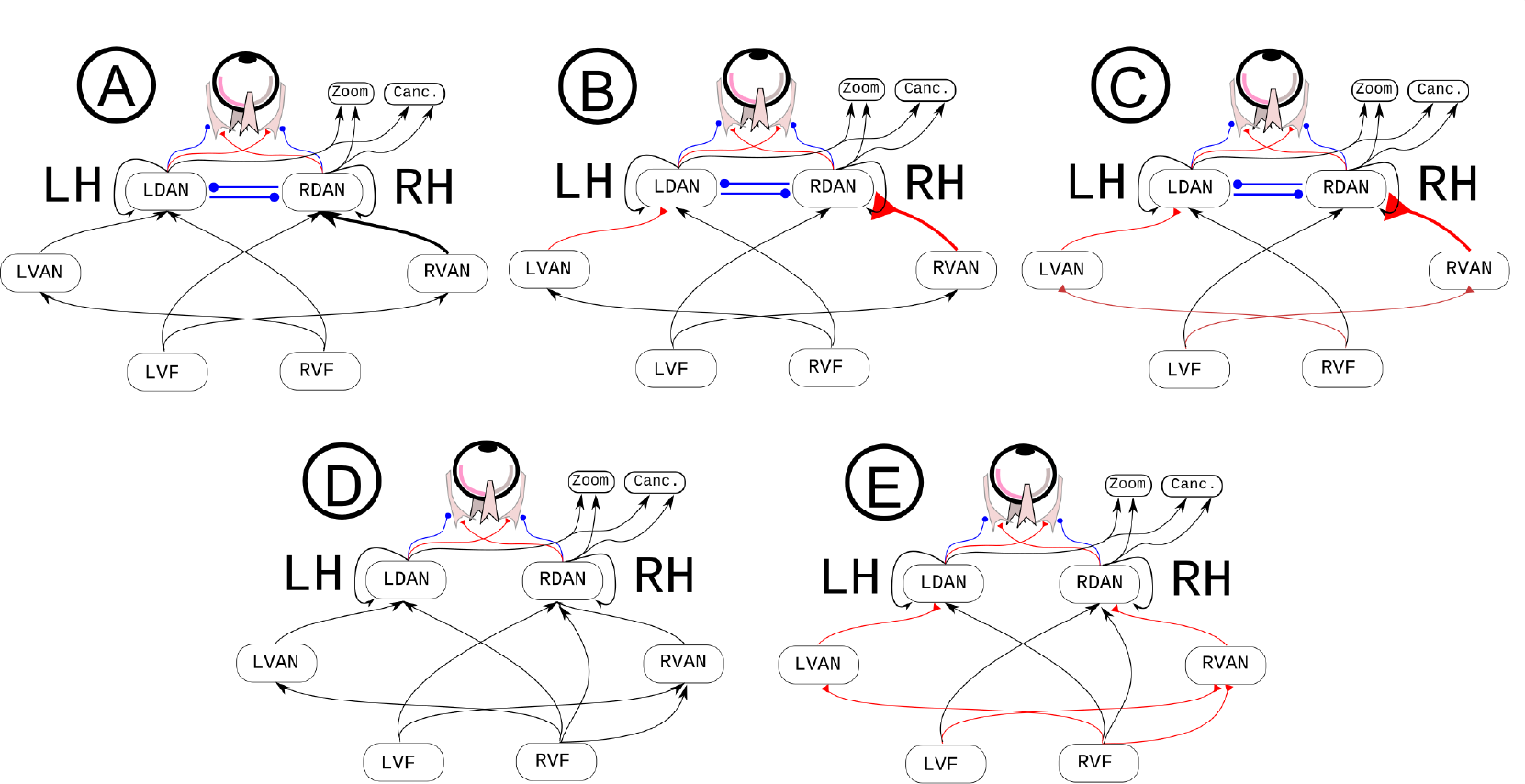
Panels A, B and C depict different implementations of the attentional networks with inter-hemispheric inhibition (Koch et al., 2011) and DAN/VAN architecture (Corbetta & Shulman, 2002). Panels D and E represent two implementations of right-hemisphere networks with bilateral competence (Heilman & Van Den Abell, 1980; Mesulam, 1981) and no inter-hemispheric inhibition. Arrows indicate connections that can be either excitatory or inhibitory; red connections with triangular arrowheads denote excitatory connections; blue round arrowheads represent inhibitory connections. LH, left hemisphere; RH, right hemisphere; Canc., cancellation units; LDAN and RDAN, dorsal attention networks in the left and in the right hemisphere, respectively; LVAN and RVAN, ventral attention networks in the left and in the right hemisphere; LVF and RVF, left and right visual field. Right and left VANs have the same number of neurons, but different patterns of connection strength.

The inter-hemispheric connections were also modeled by following anatomical and functional results obtained in the human brain, and outlined in the Introduction. Thus, (1) they connected only the DANs, but not the VANs, which thus worked in relative isolation from one another (see Fig. 9.4D in Catani & Thiebaut de Schotten, 2012) and (2) they were inhibitory, such that each DAN inhibited the contralateral one (Koch et al., 2011): each DAN induced contralaterally-directed eye movements and inhibited ipsilaterally-directed eye movements. The DANs controlled zooming and cancellation behaviors. All the hidden units within the DANs also had reentrant connections, which integrate the previous input with the current one, thus simulating a sort of simplified visual memory, in analogy to similar mechanisms occurring in the primate brain (Salazar, Dotson, Bressler, & Gray, 2012). Thus, reentrant connections resulted in some persistence of the previous inputs across steps within a given trial.

Given the importance of eye position in visually-guided target reaching (Lewis, Gaymard, & Tamargo, 1998), we provided eye position information to neurorobots through an efference copy of the motor output. In particular, motor outputs controlling the four ocular muscles were connected one to one with the four input neurons, with a fixed weight of 1 (i.e., perfect copy from input to output).

#### 3.2.2. Cancellation task

Similar to the human experiment (see section 2), neurorobots performed a 30-trial cancellation task. The human and robotic tasks were designed with the explicit constraint of being as similar as possible. Targets were presented on a virtual display measuring 512 × 512 pixels. At the start of each trial, the gaze of the artificial eye was initialized at the center of the display, with no zoom. Again, similarly to the human experiment, each trial consisted of a set of 5 round targets, with a luminance value of 0.5 (in conventional units ranging from 0 to 1.0) and a radius of 40 pixels, randomly scattered in the virtual display. Upon cancellation, targets increased their luminance to the maximum value of 1.0.

#### 3.2.3. The Adaptive/Learning process

For the present work, neurorobots were trained by means of a Genetic Algorithm, a form of evolutionary computation that implements a Darwinian process of adaptation that can model cognitive development and trial-and-error learning, especially when only distal rewards are available (Di Ferdinando, Parisi, & Bartolomeo, 2007; Nolfi, & Floreano, 2000). Genetic algorithms are a useful alternative to supervised learning in settings such as the present one, because we employed a fitness function based on the number of cancelled targets, and not a set of input-output pairings which could be used to minimize the error by a supervised learning mechanism such as back-propagation. A typical experiment starts with the generation of a random set of individual neurorobots (each defined by a specific set of parameters of a neurocontroller). Each individual is then evaluated according to a fitness function representing the desired performance on a requested task. Due to genetic operators such as mutation and crossover, the best individuals will populate the next generation. The process iterates until a specific performance or a fixed number of generations is reached. In the present work, each genetic string encodes the value of synaptic connections W_*ij*_ and neuron thresholds in the range (-5, 5). Initially, for each evolutionary experiment a set of 100 random individuals (i.e., competing sets of parameters for the neural network of the neurorobot) were generated and evaluated for their ability to find targets. Targets had to be found as fast as possible on each of 30 cancellation trials, lasting 700 time steps each. At the end of the evaluation phase, individuals were ranked according to their performance, and the best 20 were used to populate the next generation after having undergone a mutation process. Each parameter was encoded by an 8-bit string, thus mutations were implemented by bits switching with probability p=0.01. The number of generations was set to 3,000.

Three behavioral components contributed to the overall fitness, *F*: an exploration component, a component proportional to the number of target correctly cancelled, and a reward for responses promptness.

The exploration component, which was introduced to avoid the bootstrap problem (Nolfi, & Floreano, 2000), rewarded the ability of the neurorobot to explore its visual field. In particular, the area that can be explored through eye movements was split in 100 cells. Exploration fitness (*EF*) was then computed for each trial by dividing the number of visited cells by 100. A second fitness component (*TF*) was represented for each trial by the number of correctly cancelled targets divided by 5 (i.e., the total number of presented targets). Finally, a reward for promptness (*PF*) was given when all the five targets were cancelled. *PF* was inversely proportional to the number of time steps *nt*, used to cancel all the stimuli:

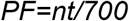

The overall fitness was calculated as

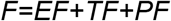

After training, neurorobots’ performance in the cancellation task was evaluated on 30 new trials, in order to measure their accuracy in finding the targets and the position of the first cancelled target, as estimated by the average value of the X coordinate of the first cancelled stimulus across trials.

#### 3.2.4. Valence of VAN-DAN connections

A set of 5 populations of neurorobots, each composed of 40 individuals, featured neurocontrollers with different connectional constraints.

Neurocontrollers A, B and C (Fig. 2) had left-right asymmetric connections between VAN and DAN (i.e., the simulated SLF II), with a greater number of connections in the right hemisphere (120) than in the left hemisphere (108). The ratio of this asymmetry difference (0.05) corresponds to the average asymmetry ratio of SLF II in 20 human subjects, as described by Thiebaut de Schotten et al. (2011) (see their supplementary Table 1). In neurocontroller A (Fig. 2A) there were no constraints in terms of type of connections (inhibitory or excitatory) along the ventral and dorsal attentional networks. In neurocontroller B a further constraint was added: VAN to DAN pathways were set to be excitatory during the training process (see Fig. 2B). Finally, in neurocontroller C also the connections projecting from the retina to the VAN were set to be excitatory (see Fig. 2C). To better evaluate the effect on performance of SLF II asymmetry, we trained two additional control populations based on neurocontroller C: C0 with completely symmetrical VAN-DAN connections (laterality ratio = 0); C1 with VAN-DAN connections only present in the right hemisphere, and absent VAN-DAN connections in the left hemisphere (complete right lateralization of SLF II).

Earlier models of spatial attention (Heilman & Van Den Abell, 1980; Mesulam, 1981) postulated a bilateral competence of the right hemisphere for both hemispaces, without explicit consideration of inter-hemispheric interactions. To simulate these models, we trained two additional populations of neurorobots (neurocontrollers D and E in Fig. 2; 40 individuals for each population). In these neurocontrollers, the right hemisphere received visual information from both the right and the left visual hemifields, while the left hemisphere received information only from the right, contralateral visual hemifield. Moreover, there were no inhibitory connections between the right DAN and its left homolog. The rest of the architecture was the same as for all the other neurocontrollers. The only difference between neurocontroller D and neurocontroller E was the valence of the connections running from the visual fields to VAN and DAN. In neurocontroller D, the valence of the visuo-attentional connections was not constrained, and could thus assume either a positive or a negative valence. In neurocontroller E, visuo-attentional connections were constrained to be excitatory, similar to neurocontroller C.

### 3.3. Results

#### 3.3.1. Behavioral Results

Figure 3 shows the ability of the five populations of neurobots to correctly solve the task. The mean percentages of correct cancellations are reported for each population.

**Figure 3.**
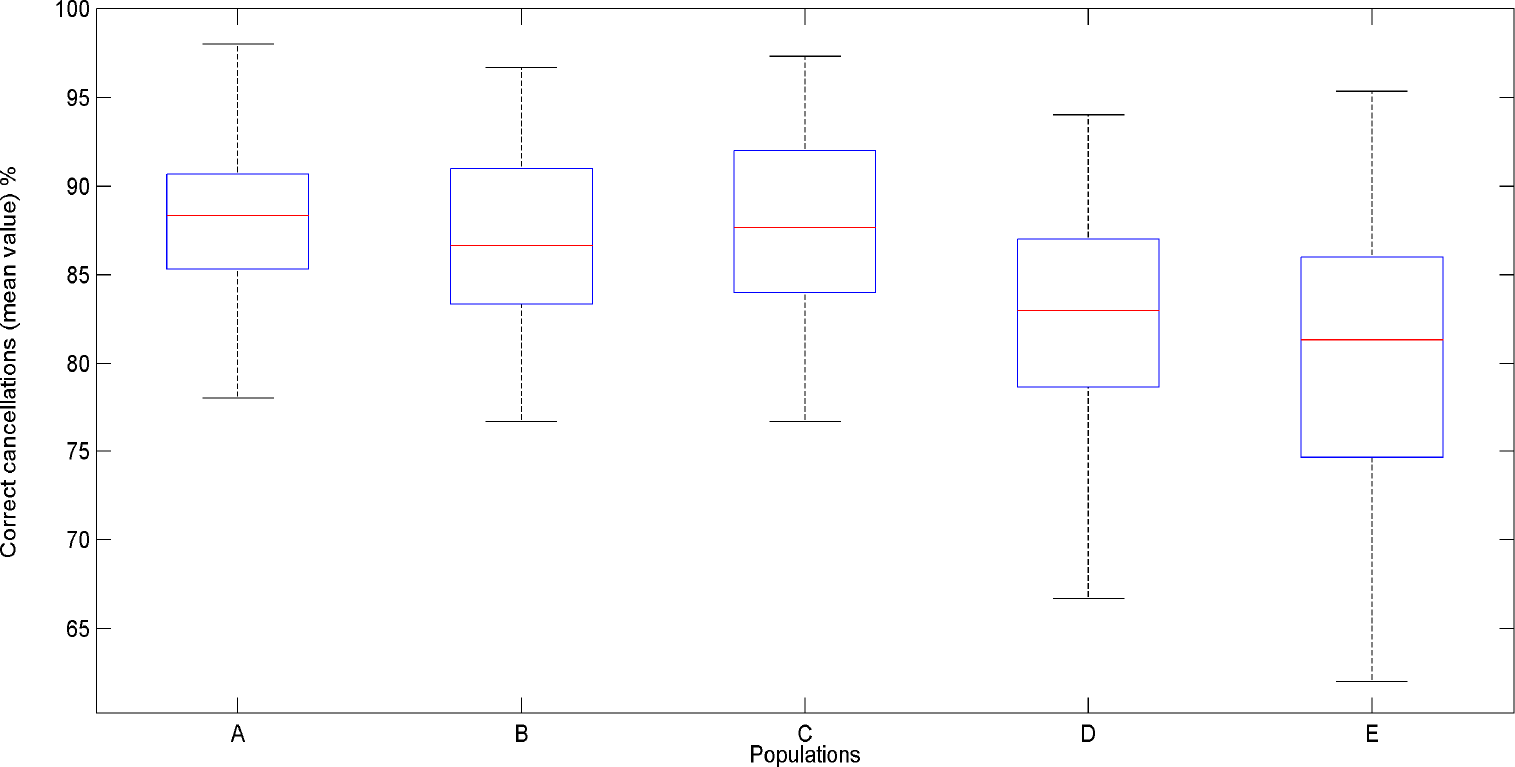
Mean percentage of correct cancellations computed across 30 trials for each population of 40 neurorobots provided with neurocontrollers A-E. The middle bar of the boxplot indicates the median of the tested population. The top and the bottom of the box indicate respectively the first (q1) and the third (q3) quartiles. Whisker length extends until the last data point that is not considered as an outlier, I.e. a point that is greater than q3 + 1.5 × (q3 – q1) or less than q1 – 1.5 × (q3 – q1). There were no outliers in the current dataset.

Figure 4 reports the performance of the three populations equipped with neurocontrollers A-E on correct cancellations. Each boxplot contains data collected for 40 neurorobots tested on 30 cancellation trials.

**Figure 4.**
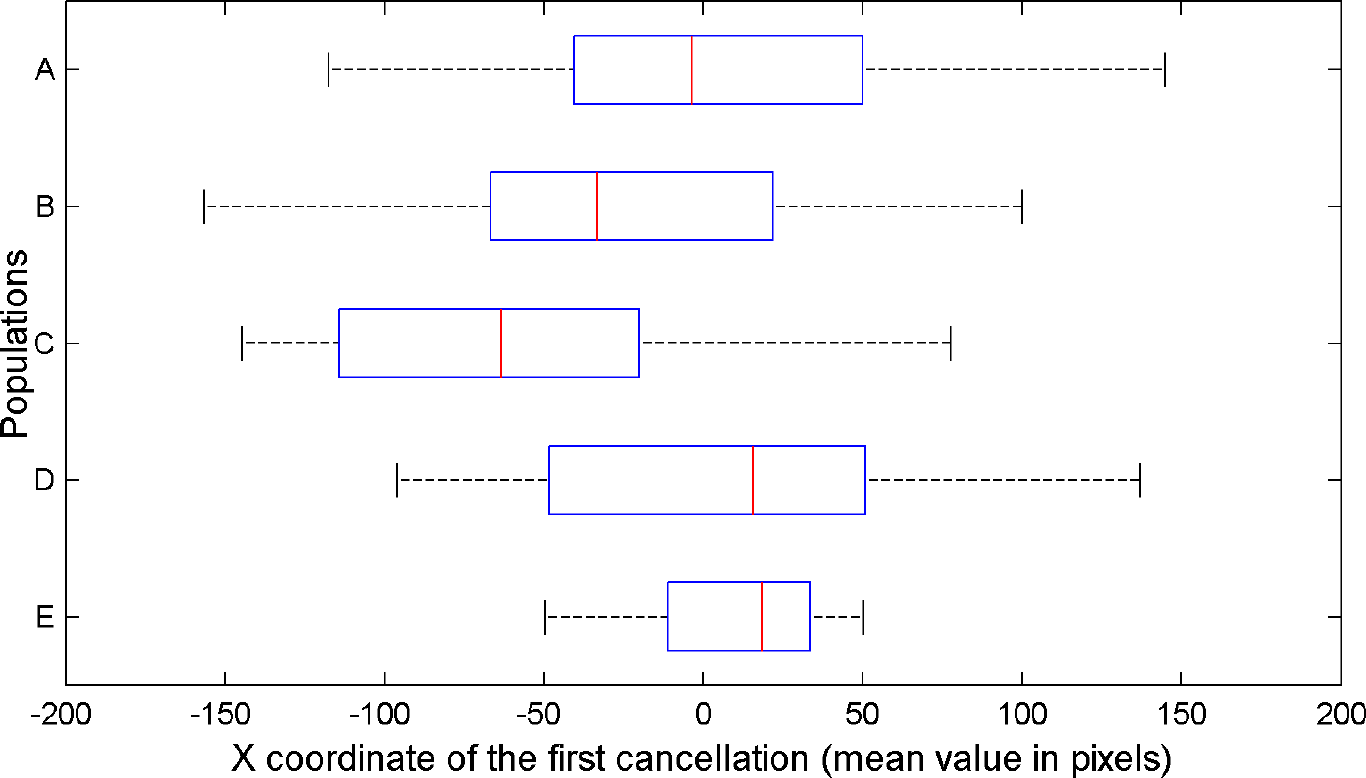
Average x values of the first cancelled target, computed across 30 trials for each population of 40 neurorobots provided with neurocontrollers A, B, C, D and E.

There were significant differences in the mean percentage of correct cancellations across the five three populations [Kruskal-Wallis test, X^2^_(4, n = 200)_ = 38.96, *p* = 7.10e-08]. Neurocontrollers with inter-hemispheric inhibition (A-C) performed better than neurocontrollers without inter-hemispheric inhibition (D-E; Post-hoc pairwise comparisons using Dunn’s-test, all ps < 0.05).

Importantly, the spatial position of the first canceled target (X coordinate value for each trial, Fig. 4) did differ across the tested populations, X^2^_(4, n = 200)_ = 34.198, *p* =4.65e-07. The position of the first canceled target was not different from 0 (central position) in neurorobots equipped with neurocontroller A (Wilcoxon-Mann-Whitney, *p*=0.1, two-tailed) and neurocontroller D (p=0.5). Neurorobots E, with bilateral competence in the right hemisphere and excitatory visual-attentional connections, showed a rightward bias, opposite to human pseudoneglect (Md=58.81, z=−2.8802, p=0.004). Neurorobots B and C tended instead to start their exploration from a left-sided target (neurocontroller B, Md = −33.27, z = −2.057, *p* = 0.02; neurocontroller C, Md = 63.29, z = −5.35, *p* < .001), thus showing a leftward bias reminiscent of human pseudoneglect. The control populations with complete SLF II symmetry (C0), or extreme rightward SLF II asymmetry (C1) showed the predicted patterns of performance: no pseudoneglect for C0 (Md=20.435, z=−0.823, *p*=0.411), and large pseudoneglect for C1 (Md=−96.526, z=−7.406, *p*=1.299e-13) (Fig. 5).

**Figure 5.**
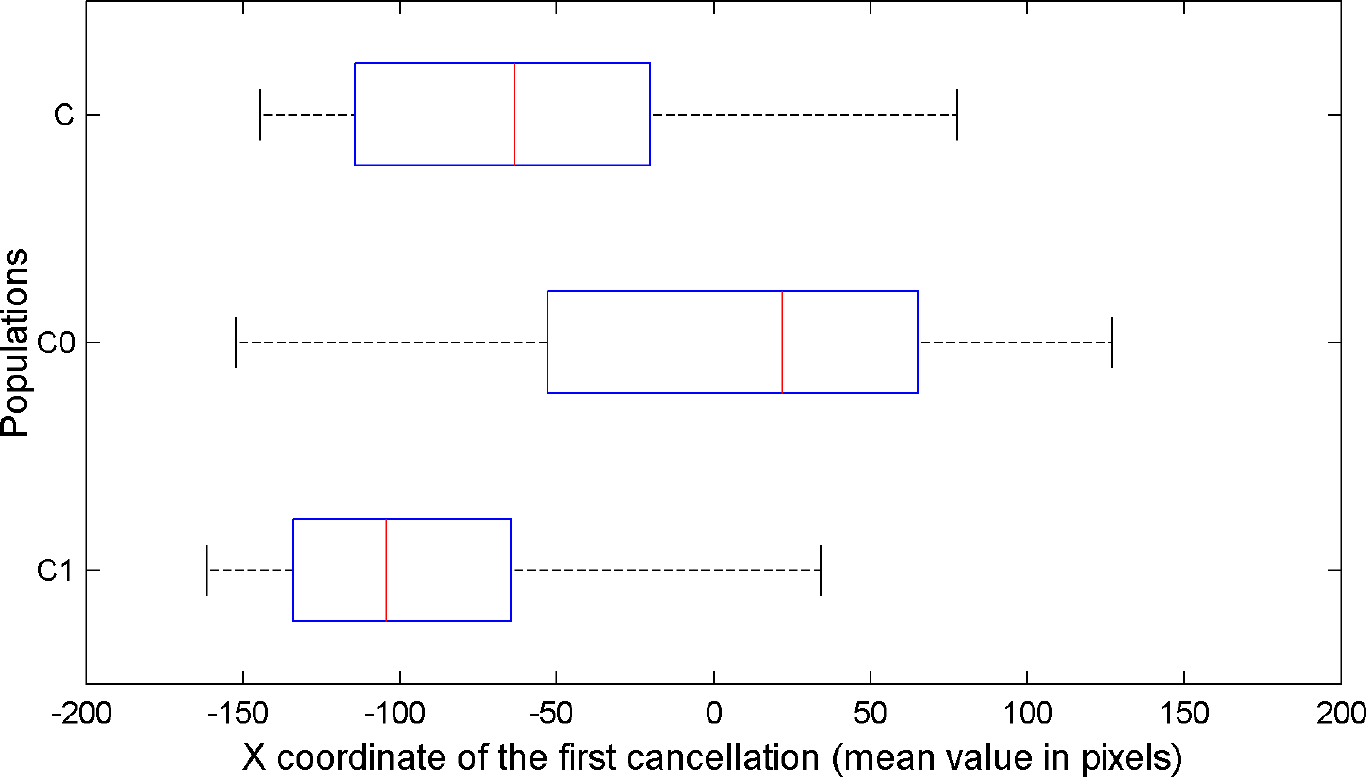
Average x values of the first cancelled targets, for all the neurorobots provided with neurocontrollers C, C0, and C1. Average x values of neurorobots C0 is not significantly different from 0, while average x values of neurocontrollers C and C1 significantly differ from 0.

**Figure 6.**
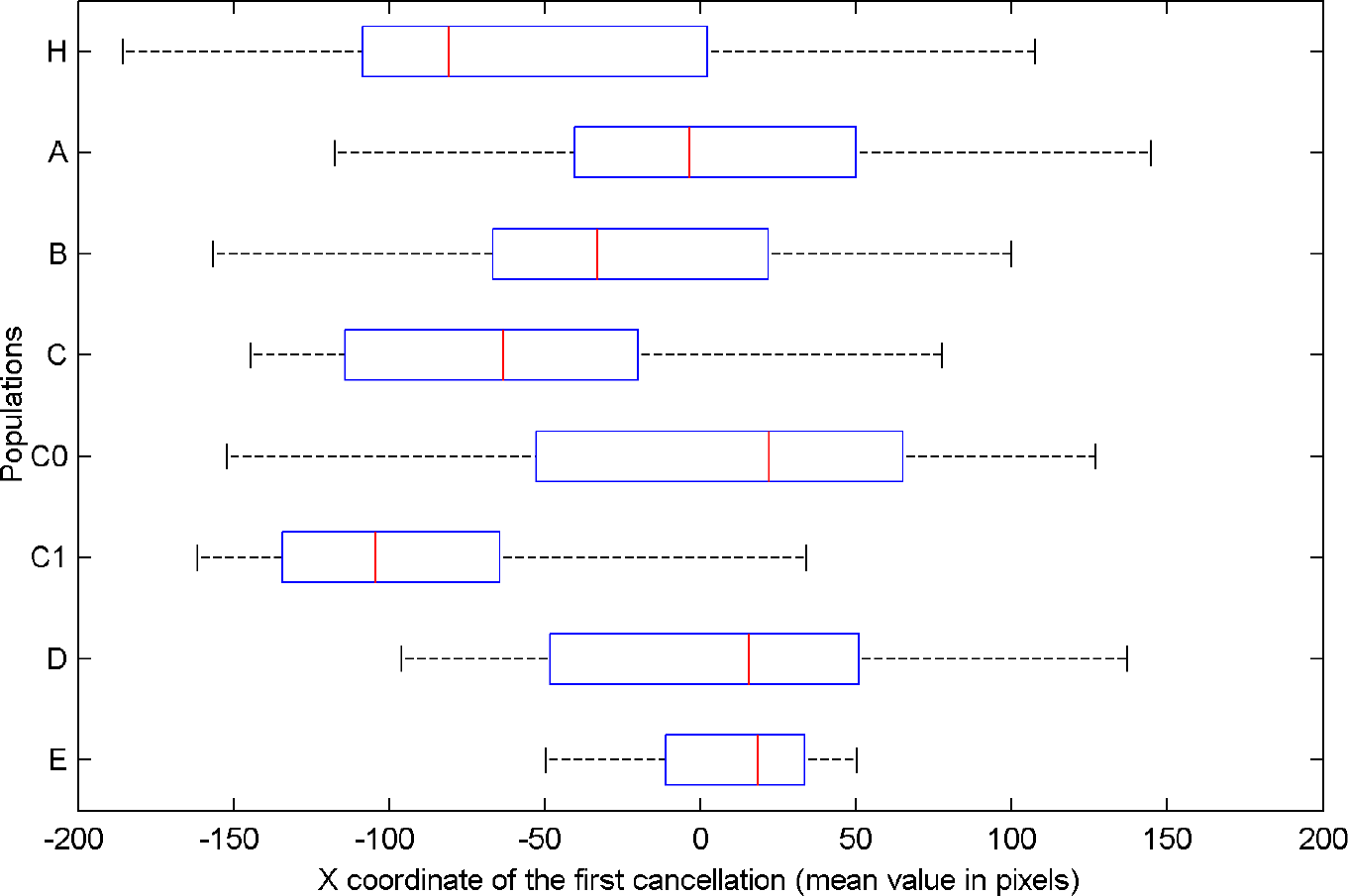
Average position on the X axis of the first cancelled targets for human participants (H) and artificial neurorobots equipped with neural networks A, B, C, C0, C1, D and E.

#### 3.3.2. Neural results

To better understand the neural dynamics leading to the exploratory bias, we examined the average activations of the DANs across all the individuals for each population, equipped with neurocontrollers C (biologically-inspired asymmetry) and C_0_ (symmetrical attention networks). We then computed a laterality index of DAN average activations between the two hemispheres: (Mean Right DAN activation - Mean Left DAN activation)/(Mean Right DAN activation + Mean Left DAN activation), with a possible range from -1 (prevalent left DAN activity) to +1 (prevalent right DAN activity). Figure 7 reports the course of the laterality index across time steps. As expected, left and right DAN activations were balanced with neurocontroller C_0_. On the other hand, in neurocontroller C activations were unbalanced toward the right hemisphere DAN. A crucial aspect for pseudoneglect concerns the initial time steps in which the exploratory bias occurs. A higher imbalance toward the right hemisphere DAN is present at the outset of the cancellation task for neurorobots C, as a consequence of asymmetries in their network architecture, while it is obviously absent for neurorobots C_0_, with symmetrical networks. The initial imbalance favoring the right hemisphere DAN is the likely basis of the spatial bias towards the initial cancellation of a left-sided item in neurorobots C.

**Figure 7.**
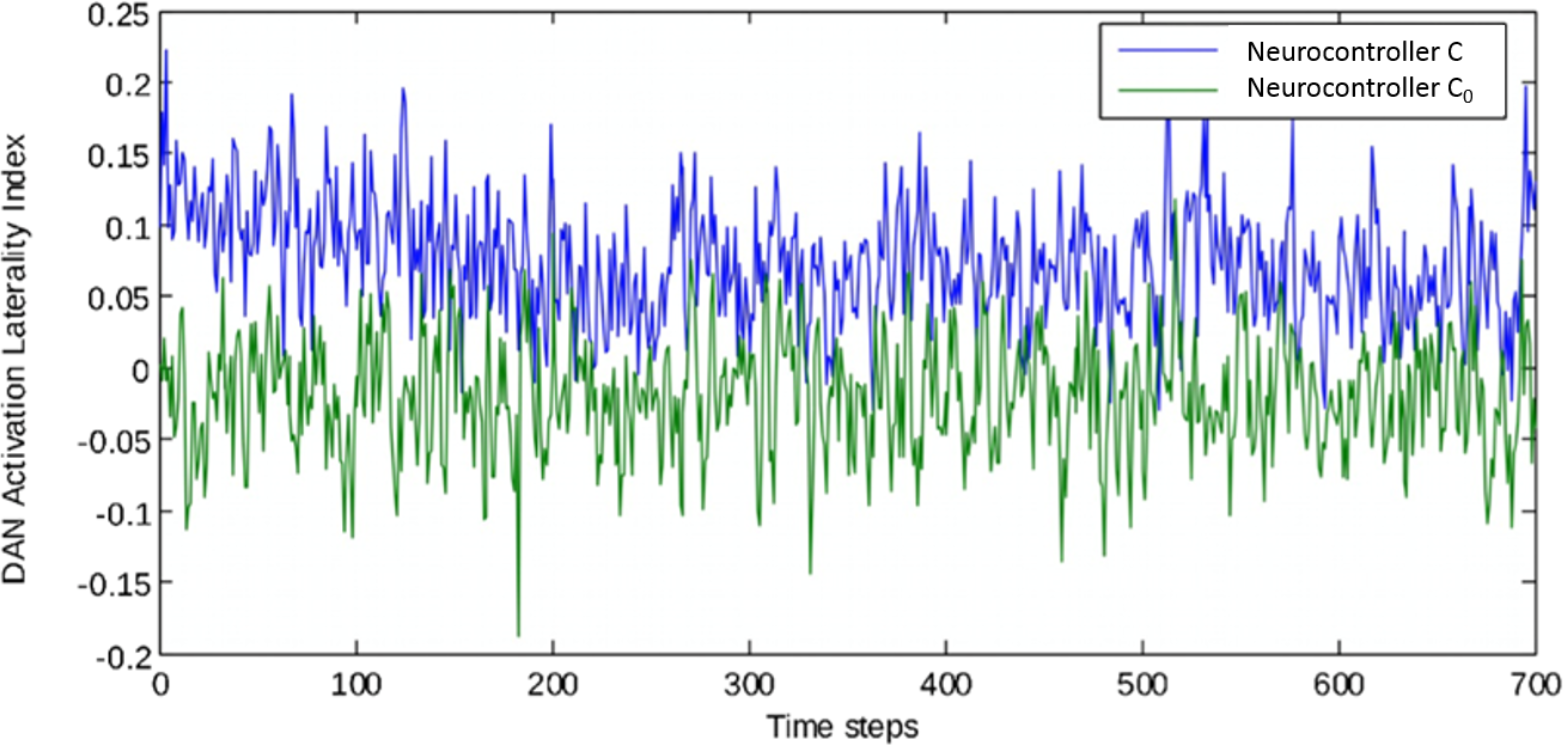
Laterality indexes of DAN activation computed for individuals equipped with neurocontroller C and C0. A value of 0 means that activation in left and right hemisphere DANs is balanced; positive values denote prevalence of right hemisphere DAN, negative values indicate prevalence of left hemisphere DAN.

Figure 8 shows the average activation of the hidden DAN neurons in the left and in the right hemisphere during the first 30 time steps of the cancellation task, for agents equipped with the biologically inspired neurocontroller C, and for those equipped with the symmetrical neurocontroller C_0_. The initial activation is symmetrical for the C_0_ agents, but it is higher in the right hemisphere than in the left hemisphere for the C agents. Thus, an asymmetry of VAN connections results in a corresponding activation asymmetry in the anatomically symmetrical DANs. The DAN asymmetry in the initial phases of the task is the simulated neural correlate of behavioral pseudoneglect. After the initial phase, the left-right differences are absorbed by the increased activity of the hidden units; when left and right activities reach saturation, the behavioral asymmetry decreases (see Fig. 7, where asymmetry of performance decreases around time step 150 for neurocontroller C).

**Figure 8.**
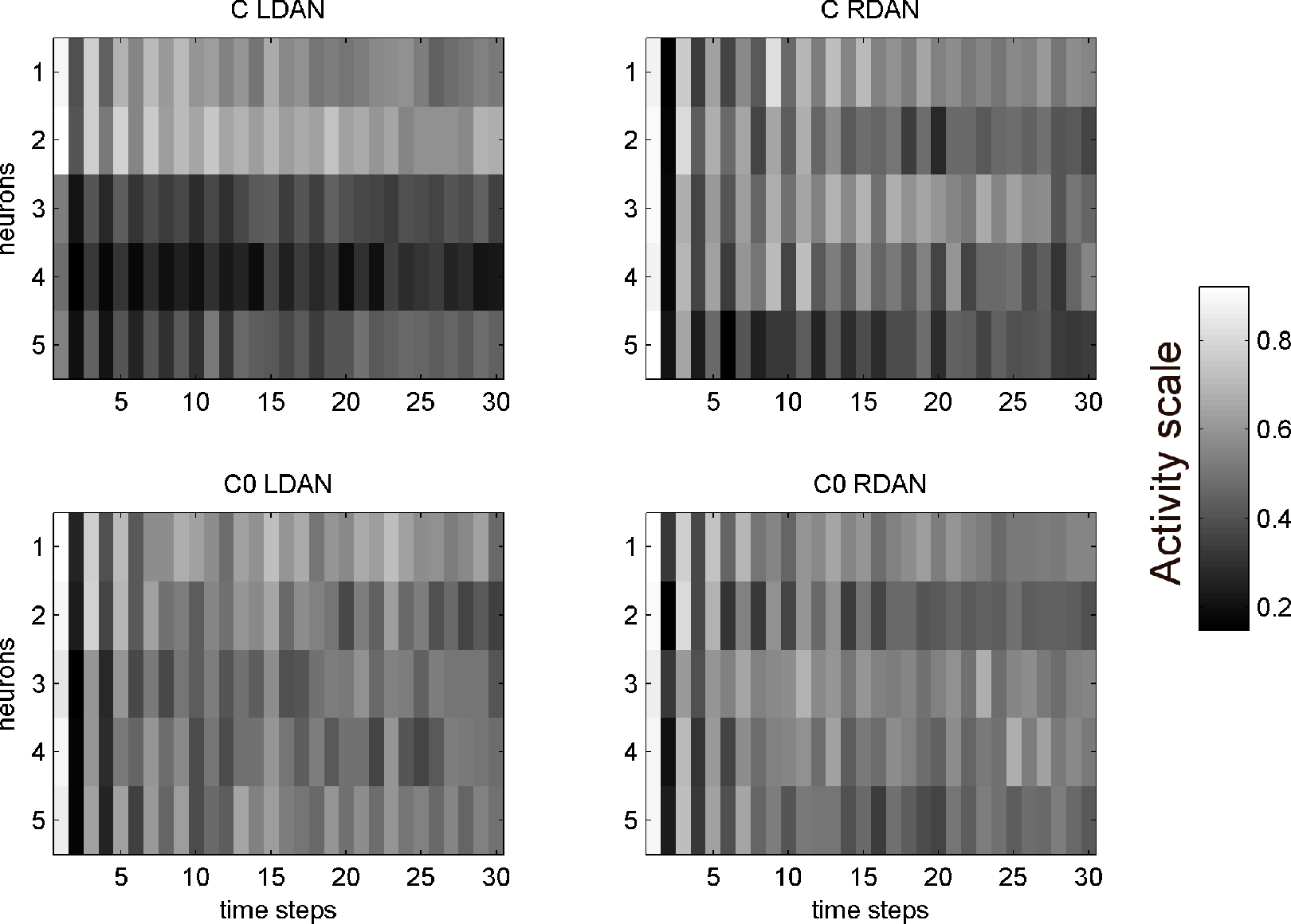
Average activation of hidden neurons in right hemisphere DAN (RDAN) and in left hemisphere DAN (LDAN), for the first 30 steps of individuals equipped with neurocontrollers C and C0. The activity scale goes from 0 (black) to 1 (white). Note the early, large left-right asymmetry in neurobiologically inspired C agents (arrows), which subsequently decreases. The symmetrical C0 agents do not show any asymmetry of performance.

#### 3.3.3. Comparison between human and robotic performance

Human participants and robotic populations as a whole did not show the same distribution of the position of the first cancelled targets (Kruskal-Wallis test, X^2^(5, n = 301) = 67.88, *p* < .001) (see Fig. 6). Post-hoc tests (Dunn’s test with Bonferroni correction) demonstrated a difference in distribution between humans and neurocontrollers A (*p* <.001), B (*p*=0.0394), C_0_ (*p* < .001), C_1_ (*p* = 0.0153). However, the position distribution derived from human performance and neurocontroller C’s performance showed a similar degree of leftward asymmetry (Fig. 9; Dunn’s test, *p* = 1.0; Levene test of homogeneity, *p* = 0.39). Thus, all robotics agents performed differently from humans, with the notable exception of the neurorobot population C, whose performance provided a good approximation to human performance.

**Figure 9.**
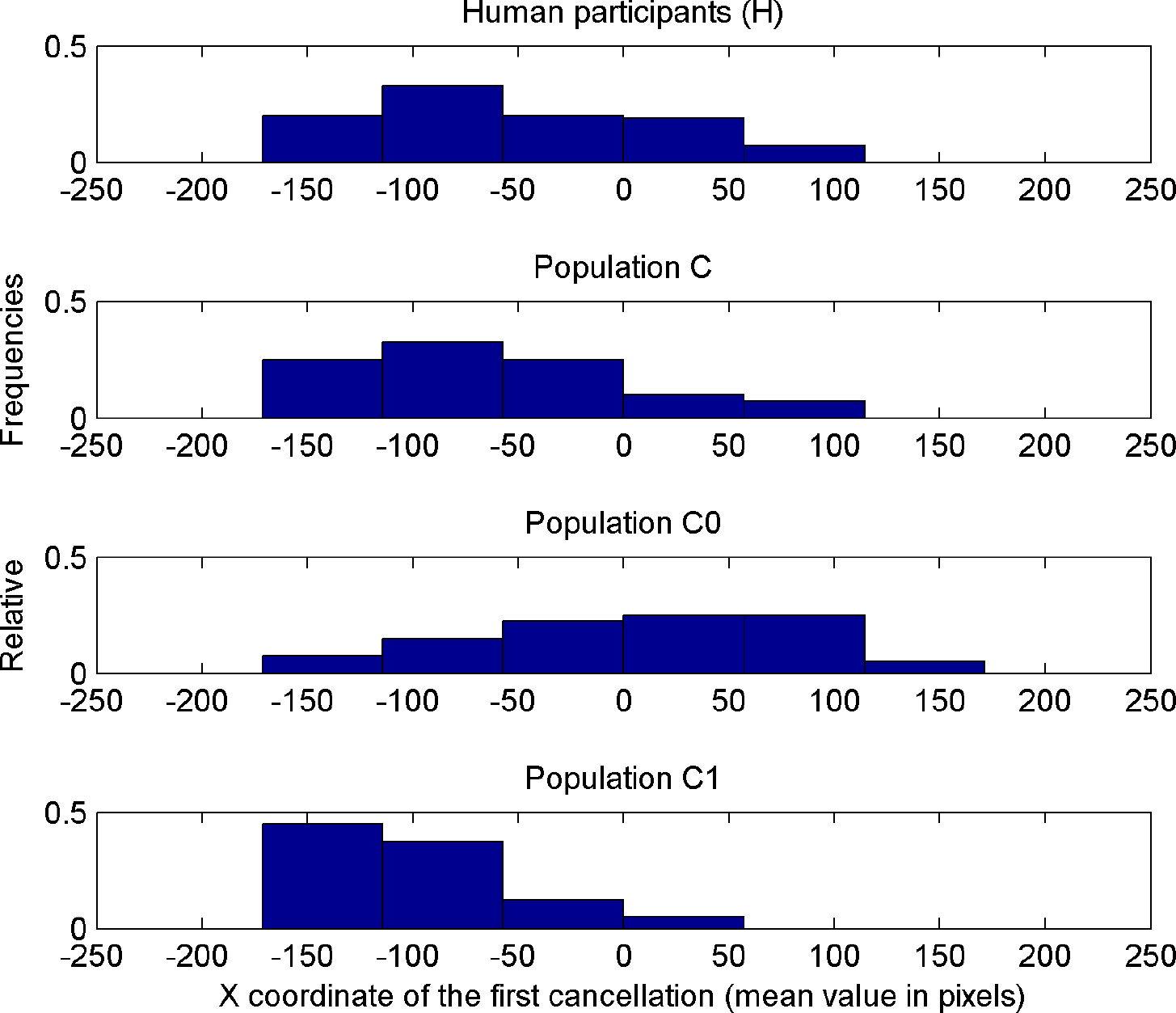
Relative frequencies of the distribution of the position of the first cancelled target for 101 human participants (see Experiment 1) and for the populations of neurorobots C (equipped with the biologically inspired neurocontroller), C0 (presenting symmetrical DAN) and C1 (with VAN-DAN connections only present in the right hemisphere).

We then compared the performance over time of human participants and model C neurorobots not only for the first canceled target (see Fig. 9), but across all the presented targets. We performed a Bayesian repeated measures ANOVA (JASP software, version 0.8.2), with agents (human, neurorobots C) as between-group factor, and the spatial position (X coordinate) of the sequence of all the five canceled targets as within-group factors. The Inclusion Bayes Factor, which compares ANOVA models that contain a given effect to equivalent models stripped of the effect, showed decisive evidence (BF *Inclusion*= 2.137e +42) for the cancellation order main effect. Thus, the order of cancellation of all the five targets depended on their spatial position (Fig. 10). Importantly, this effect was statistically equivalent for the human and the neurorobot C populations. In particular, there was substantial evidence against the existence of a group main effect (BF*Inclusion* = 0.144), and strong evidence against the existence of a group X cancellation-order interaction (BF*Inclusion* = 0.046). These results show that the neurorobots from population C and human subjects behave similarly over time when canceling all the five targets.

**Fig. 10.**
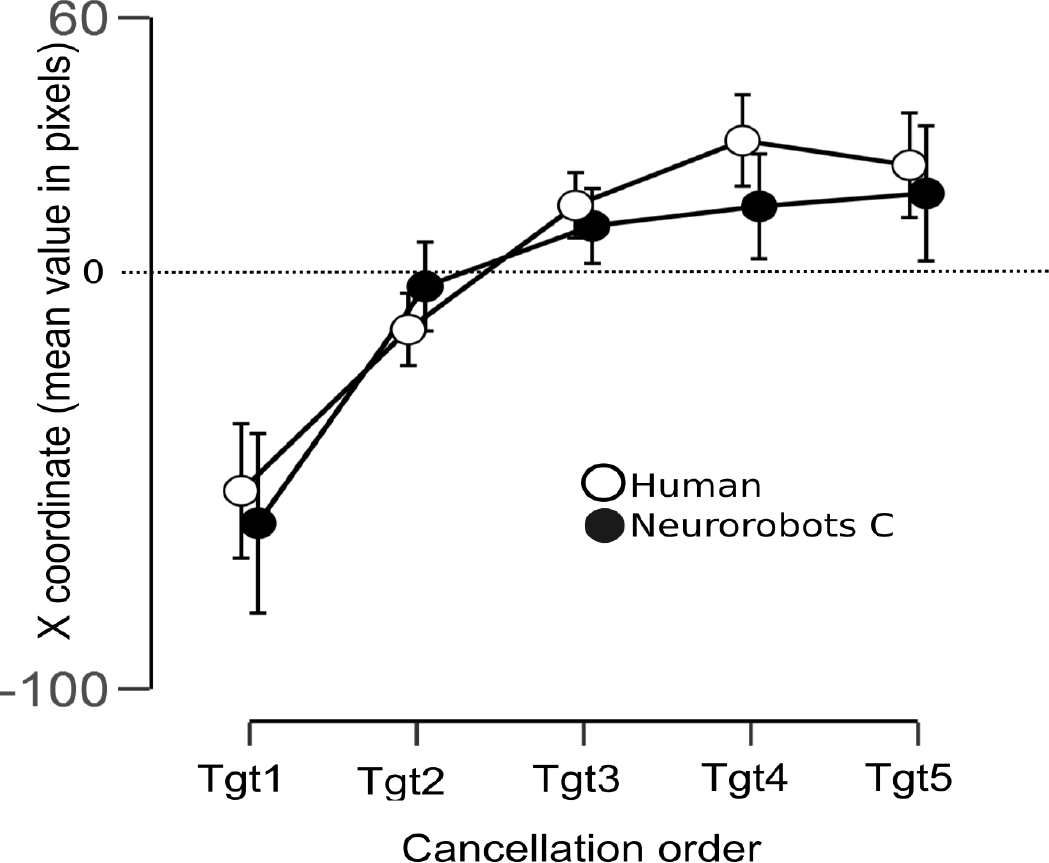
Coordinates of canceled targets as a function of the temporal sequence of cancellation in human participants and in neurorobot population C. Error bars represent credible interval of 95%

## 4. General Discussion

In this study, we established specific connectivity constraints leading to a lateral spatial bias (pseudoneglect) in artificial organisms trained to perform a visual search task by using genetic algorithms. A form of pseudoneglect that was qualitatively and quantitatively similar to that shown by normal participants did emerge in artificial neurorobots, but only in those harboring hemispheric asymmetries of connectivity that simulated those typically occurring in the human brain. As a further condition, a general excitatory influence of VAN on the ipsilateral DAN was necessary for pseudoneglect to occur in neurorobots. This novel result suggests that hemispheric asymmetry alone is not sufficient to generate a leftward bias, and thus further specifies the likely connectional constraints of pseudoneglect.

We first consider our results in the light of neurophysiological studies of pseudoneglect, and then in relation to existing modeling studies of the human attentional system. A particular instance of pseudoneglect occurs with the landmark task: When judging lines pre-bisected to the left of their true center, normal participants consider the left segment as being longer than the right one (Milner, Brechmann, & Pagliarini, 1992). Spatial attention has been shown to be a major determinant of this phenomenon (Toba et al., 2011). Szczepanski et al. (2013; 2010) tested normal participants’ spatial bias on convert attention tasks and on the landmark task by using a multimodal approach, combining psychophysics, fMRI and TMS. They tested only frontal and parietal ROIs in the DAN, and did not explore the VAN. Their subjects’ sample showed a mixed spatial bias: some subjects had a leftward bias (pseudoneglect), but most subjects showed a rightward bias (Szczepanski & Kastner, 2013). On average, the bias was rightward, unlike most of the literature results. The lateralization of the bias correlated with the lateralization index of the fMRI activation in the ensemble of the DAN ROIs during a covert spatial attention task. Specifically, subjects that had more left hemisphere activation also had a contralateral, i.e. rightward, bias in the landmark task; conversely, subjects with more right hemisphere activation tended to have a leftward behavioral bias. TMS-induced interference on the left-or right-hemisphere parietal nodes during the landmark task caused an ipsilateral shift of the bias: right parietal TMS caused a rightward shift compared to the initial bias, and left parietal stimulation caused a leftward shift. Stimulating both right and left parietal ROIs did not cause a shift. Szczepanski and Kastner (2013) suggested that there is an inter-hemispheric competition between the DAN nodes, and the lateralization of the sum of the weights in the DAN activation shifts the attentional focus contralaterally. Thus, these results are broadly consistent with the functioning of the present neurorobot population C. In agreement with Szczepanski and Kastner’s (2013) conclusions, the DAN in the current model is conceptualized as a whole, and not as separated nodes. Additionally, Szczepanski and Kastner’s data showed that there is large variability between participants in the direction and degree of lateralization of DAN activation, that on average did not significantly differ between the hemispheres. Here we aimed to explore the typical functional architecture in the human population. Therefore, we chose to model the DAN as laterally symmetrical and the VAN as right-lateralized. However, there are several differences between the current models and the Szczepanski et al’s studies. First, they used a landmark task while here we used a search task. Second, the overall behavioral pattern here was of a leftward classical pseudoneglect bias and not the rightward bias found by Szczepanski et al. This might result from substantial differences in the studied samples or in the tasks used. Third, and more importantly, the VAN, which has a major contribution in the current model, was not tested in their studies.

The architecture of neurorobot C is partly inspired by the results of Koch et al (2011), which might oversimplify the nature of interhemispheric interactions. Several fMRI studies of human attention areas found evidence of bilateral activation of attention areas, with a contralateral bias (see, e.g., Patel et al., 2015). In neurorobots D and E, we introduced bilateral competence in the right hemisphere networks (Heilman & Van Den Abell, 1980; Mesulam, 1981). However, performance this model showed no consistent spatial bias. This suggests that right hemisphere bilateral competence by itself might not be crucial to the emergence of pseudoneglect. On the other hand, it is true that evidence from neglect patients (Bartolomeo & Chokron, 1999) challenges models of attention exclusively based on inter-hemispheric rivalry (Kinsbourne, 1970, 1977, 1993), and that bilateral competence in attentional areas might be important in long-term compensation of neglect (Bartolomeo & Thiebaut de Schotten, 2016; Lunven et al., 2015). Our results stressing the importance of both right-hemisphere bilateral competence and inter-hemispheric competition for pseudoneglect may thus pave the way for an integrated interpretation of different lines of research on normal or dysfunctional human attention networks.

In their recent review, Borji and Itti (2013) provided a taxonomy of nearly 65 computational models of visual attention. Many of these models focused on reproducing eye movements [e.g., the saliency-based models reported in Borji and Itti (2013)], following a bottom up approach. Typically, these models extract a set of features, represented as maps, from an incoming image. Then, feature maps are combined in a saliency map where a winner-take-all mechanism will designate the spatial region to be attended. Saliency-based attention models in general do not account for exploration biases, with the exception of a recent model (Ali Borji & Tanner, 2016), where an object center bias (the tendency to focus on the center of objects) is reproduced by adding an ad-hoc bias map to the saliency map. While important for building predictive models, this result seems little relevant to lateral biases such as pseudoneglect. Other models (Deco & Rolls, 2004; Deco & Zihl, 2004) simulate attention as emerging from the competition of several brain areas subjected to bottom-up and top-down biases. These models do not drive eye movements; the scan path is simulated as a sequence of activations of the simulated posterior parietal cortex. Lanyon and Denham (2004, 2010) added to these models simulated eye movements and an adjustable attention window scaled according to stimuli density. Despite being successful at reproducing scan paths in healthy individuals and neglect patients, these models do not address the issue of pseudoneglect. Other models of attention did not consider pseudoneglect because of their training procedure or design constraints (Di Ferdinando et al., 2007; Monaghan & Shillcock, 2004; Mozer, 2002; Pouget & Sejnowski, 2001). Di Ferdinando et al. (2005) explored line bisection and target cancellation performance in four biologically inspired neural networks. The networks’ patterns of connectivity varied along different degrees of asymmetry, inspired by specific theories. Pseudoneglect occurred in line bisection but not in visual search. In these models, motor outputs were only used for target selection; there was no active exploration of the environment, whereas when our neurorobots explored their environment the corresponding input information changed as a function of eye movements. Nonetheless, the present study shares with Di Ferdinando et al. (2005) and other work from the Zorzi group (Casarotti, Lisi, Umiltà, & Zorzi, 2012) the stress on accounts of attentional phenomena relying on sensory-motor transformations, as stated by the premotor theory of attention (Rizzolatti, Riggio, Dascola, & Umilta, 1987). Specifically, our results support the hypothesis that the way in which the movements of the actuators are controlled affects the performance on a cancellation task (Gigliotta, Bartolomeo, & Miglino, 2015).

Thus, contrary to most available models of attention, our artificial robots are trained to correctly cancel target stimuli, and are free to self-organize in order to find a proper solution, within the sole limits of the imposed connectivity constraints. These constraints were inspired by available data concerning the anatomical and functional organization of the attentional networks in the human brain. To the best of our knowledge, this is the first attempt to simulate the dorsal and ventral attention networks in the two hemispheres of the human brain. Another original feature of the present models is the embodiment factor, consisting of the explicit modeling of eye and hand movements (see also Bartolomeo, Pagliarini, & Parisi, 2002; Di Ferdinando et al., 2007; Gigliotta et al., 2015; Lanyon & Denham, 2004; Miglino, Ponticorvo, & Bartolomeo, 2009). In particular, the present models extended the models devised by Di Ferdinando et al. (2007), by increasing the complexity of the organisms’ retina, the biological plausibility of the motor system and that of the neural controllers. Conti et al. (2016) also adopted an embodied perspective, based on a humanoid robot platform. In their study, an iCub robot was trained to remove objects from a table, a task reminiscent of a cancellation task. Intra-hemispheric disconnections were able to produce neglect-like behavior. However, the embodiment of the model was limited by the facts that selection of a visual target was carried out independently of the motor behavior, and that robot’s eyes were kept fixed during the cancellation task. Moreover, although hemisphere asymmetry was modeled by increasing the number of right hemisphere processing units, no bias in normal performance is reported.

Moreover, contrary to most published work, our model attempted to simulate the relationships between the visual pathways and the attentional networks by respecting important biological constraints. Visual pathways project mainly to the hemisphere contralateral to each visual field. However, theoretical models of visual attention posit that the left hemisphere mainly deals with the contralateral hemispace, whereas the right hemisphere has a more bilateral competence (Heilman & Van Den Abell, 1980; Mesulam, 1981). In previous computational models this asymmetry has not always been simulated in a biologically plausible way. In some cases, both simulated hemispheres received visual information from the whole visual field, with attention asymmetries being represented in inner layers (Di Ferdinando et al., 2007; Monaghan & Shillcock, 2004). In the Conti et al.’s model (Conti et al., 2016), the right hemisphere received information from both visual hemifields, whereas the left hemisphere processes only the contralateral visual hemifield. Our models D and E had similar architecture, but were unable to mimic human performance. Moreover, there is no anatomical evidence of such asymmetries in the visual pathways, and information exchange in the occipital visual areas is mainly limited to the vertical meridian (Berlucchi, 2014). In our model, these important biological constraints of visual information processing were respected, because each artificial hemisphere received visual information from the contralateral hemifield; inter-hemispheric connections were only present at a later stage of processing, between the artificial DANs.

It might be argued that in our model C a leftward bias was simply transferred or amplified from the input to the output layers. If so, however, we would have expected to observe a constant leftward bias, akin to right-sided neglect. What we found, instead, was just an initial leftward bias, at the onset of the exploration task, analogous to human physiological pseudoneglect. In order to observe this initial bias, the VAN-DAN connections had to have an excitatory valence. This occurrence does not result from existing empirical data and is thus a novel prediction of the model. Also, neurorobot populations D and E, which also had more right hemisphere than left hemisphere resources, and should then entail a similar input-to-output amplification, did not show pseudoneglect, presumably because of the lack of inter-hemispheric inhibition.

The level of detail of the models is not a trivial matter, because it has to provide meaningful novel information while remaining tractable. A potential limitation of our study is the use of simplified versions of the fronto-parietal cortical networks, without taking into consideration the substructures of the DAN and VAN, which are both broad and partly heterogeneous networks (Colby & Goldberg, 1999), nor subcortical structures such as striatum, thalamus and superior colliculus (Krauzlis, Bogadhi, Herman, & Bollimunta, 2017). For example, the connectional anatomy of VAN components such as the temporo-parietal junction (e.g., with the ventral cortical visual stream) and of the ventro-lateral prefrontal cortex (e.g., with limbic structures) is likely to be crucial to the functioning of the VAN. Yet, our simplified model, with a VAN receiving visual input and sending excitatory connections to the ipsilateral DAN, was able to mimic human performance to an impressive level of accuracy.

More generally, our modeling is consistent with evidence from healthy subjects and neglect patients, stressing the importance of entire fronto-parietal networks, or of their dysfunction, in behavioral patterns such as pseudoneglect (Szczepanski & Kastner, 2013), or visual neglect (Bartolomeo et al., 2012; Corbetta & Shulman, 2011), respectively. Also, integrated fronto-parietal activity, with subtle, task-dependent differences in network dynamics, occurs during attention orienting in monkeys (Buschman & Miller, 2007). Concerning visual neglect, evidence suggests that a major determinant of this condition is indeed a dysfunction of the right hemisphere VAN (Corbetta & Shulman, 2011; Urbanski et al., 2011), or of its connections with the ipsilateral DAN (Thiebaut de Schotten et al., 2005).

Finally, we note that the present population-based model can be potentially used to explore in a natural manner the universal properties (the basic brain architecture) and individual differences in network efficiency, two aspects recently underlined by Michael Posner (2014) as appropriate features for future models of attention.

In conclusion, we have demonstrated the emergence of pseudoneglect behavior in artificially evolving neurorobots searching for visual objects, under specific connectional constraints. These neurorobots provide a plausible model for the dynamic interactions between fronto-parietal attention networks in the human brain.

